# Establishment of *Etv5* gene knockout mice as a recipient model for spermatogonial stem cell transplantation

**DOI:** 10.1101/2020.03.10.985408

**Authors:** Xianyu Zhang, Xin Zhao, Guoling Li, Mao Zhang, Pingping Xing, Zicong Li, Bin Chen, Huaqiang Yang, Zhenfang Wu

**Author notes:** Correspondence: Huaqiang Yang,; Bin Chen,; and Zhenfang Wu.

## Abstract

Transplantation of spermatogonial stem cells (SSCs) is an alternative reproductive method to achieve conservation and production of elite animals in livestock production. Creating a recipient animal without endogenous germ cells is important for effective SSC transplantation. However, natural mutants with depletion of SSCs are difficult to obtain, and drug ablation of endogenous germ cells is arduous to perform for practical use. In this study, we used mouse models to study the preparation of recipients with congenital germ cell ablation. We knocked out (KO) Ets-variant gene 5 (*Etv5*) in mice using the CRISPR/Cas9 system. The testicular weight of *Etv5*^-/-^ mice was significantly lower than that of wild-type (WT) mice. The germ cell layer of the seminiferous tubules gradually receded with age in *Etv5*^-/-^ mice. At 12 weeks of age, the tubules of *Etv5*^-/-^ mice lacked germ cells (Sertoli cell-only syndrome), and sperm were completely absent in the epididymis. We subsequently transplanted allogeneic SSCs with enhanced green fluorescent protein (EGFP) into 3-(immature) or 7-week-old (mature) *Etv5*^-/-^ mice. Restoration of germ cell layers in the seminiferous tubules and spermatogenesis was observed in all immature testes but not in mature adult testes at 2 months post-transplantation. The presence of heterologous genes *Etv5* and *EGFP* in recipient testicular tissue and epididymal sperm by PCR indicated that sperm originated from the transplanted donor cells. Our study demonstrates that, although *Etv5*^-/-^ mice could accommodate and support foreign germ cell transplantation, this process occurs in a quite low efficiency to support a full spermatogenesis of transplanted SSCs. However, using *Etv5*^-/-^ mice as a recipient model for SSC transplantation is feasible, and still needs further investigation to establish an optimized transplantation process.

## Introduction

Spermatogonial stem cells (SSCs) are male germline stem cells that reside in the basement membrane of the seminiferous tubule in the testis (Scadden, 2006). SSCs are capable of self-renewing themselves to maintain the stem cell pool throughout the lifespan and differentiating into spermatozoa after puberty (Yoshida, 2012). SSCs form the foundation of spermatogenesis and male fertility. SSCs share some common identities with other adult stem cells, but they also harbor unique and important function by transmitting genetic information from the paternal generation to the descendants (Komeya., 2015). SSCs can be isolated, propagated in vitro, cryopreserved, and transplanted into the recipient testis to generate SSC-derived progeny (Kubota., 2018). The potential to manipulate or transplant SSC has offered a new approach to repopulate sterile testis and restore spermatogenesis in animal models or patients. In 1994, Brinster and colleagues first reported that transplantation of mouse SSCs into the seminiferous tubules of infertile recipient mice reinitiates donor-derived spermatogenesis to produce viable offspring (Brinster., 1994). Since then, SSC transplantation has also been demonstrated in many species, including rat, goat, sheep, pig, and even non-human primate (Ogawa., 1999; Honaramooz., 2003; Rodriguez-Sosa., 2009; Mikkola., 2006; Hermann., 2012). This technique opens new avenues for the treatment of male infertility, development of alternative livestock reproduction technology, and generation of transgenic animals for biomedical and agricultural purposes.

SSC transplantation is routinely performed in both mice and rats but has not yet been well-established in large mammals, such as pigs and cattle (Mikkola., 2006; Herrid., 2006). SSC transplantation in large farm animals is a promising technique for fast multiplication of elite or genetically desired individuals to benefit agricultural outputs (Giassetti., 2019). In addition, translation of this technology into human clinics has not been realized (Nagamatsu., 2017). Full implementation of the transplant relies largely on effective derivations of germline-ablated recipient and high-quality donor cells. Various approaches, such as drug treatment (Brinster., 1994), irradiation (Herrid., 2009), and heat shock (Ma., 2011), have been adopted to generate a sterile recipient for SSC transplantation. The most common and effective strategies for germ cell ablation in mouse models are injection of chemotoxic drugs (busulfan) and use of mutant W mice lacking endogenous germ cells. However, these strategies are not easily reproduced in large animals, as an optimal busulfan dose to balance an adequate ablation of endogenous germ cells and whole body side effect is difficult to control; moreover, no other species has a W genetic background which remains unclear for its mutation information (Kubota., 2018). The issue of recipient preparation prompted us to develop a common and simple method to generate germline-ablated recipients. This model should feature (i) congenital germ cell deficiency; (ii) intact Sertoli cells and testicular structure to support foreign SSC residence, growth, and differentiation; (iii) transmissibility of phenotype to progeny; and (iv) easy application to various animal species. On the basis of the above considerations, specific gene mutations associated with the developmental deficit of SSCs but unaffecting testicular somatic support cells can be used to create such a recipient model. In this regard, the currently developed genome editing technology can engineer any genomic regions to generate desired genotypes and phenotypes in many species, therefore offering a universal platform for genome engineering in various animals.

Here, we establish an Ets-variant gene 5 (*Etv5*) gene-targeted mouse model and investigate its ability to support allotransplantation of SSCs. Genetic variants in the human *Etv5* gene was believed to be associated with nonobstructive azoospermia associated with Sertoli cell-only syndrome (O’Bryan., 2012). *Etv5* homozygous mutant (*Etv5*^*-/-*^) male mice are sterile due to the progressive loss of germ cells. *Etv5* heterozygous mutant (*Etv5*^*+/-*^) mice are fertile and healthy, so they can be used as parental generation to maintain the mutant strain. Our study explored the possibility to use *Etv5*^*-/-*^ mice as the recipient model with congenital germline ablation to facilitate SSC transplantation study. If *Etv5*^*-/-*^ mice is applicable for SSC transplantation, this strategy can be extended to large farm animals by specific gene targeting to create transplant recipients serving SSC-based reproduction and transgenesis.

## Materials and methods

### Ethics statement

All animal experiments were performed in accordance with the guidelines of the National Institutes of Health’s Guide for the Care and Use of Laboratory Animals. Our study was approved by the IACUC at South China Agricultural University.

### Embryo injections for KO mice production

CRISPR-gRNA targeted sequences are shown in Fig. 1A. The mouse (C57BL/6) *Etv5* gene (GenBank accession number: NM_023794.2; Ensembl: ENSMUSG00000013089) is located on chromosome 16, with 13 identifiable exons, with an ATG start codon on exon 2 and a TAA stop codon on exon 13. The introns on both sides of exons 1–5 were selected as the target sites for CRISPR/Cas9-mediated genome editing. A pair of gRNA targeting vectors (pRP[CRISPR]-hCas9-U6) was constructed and confirmed by sequencing: gRNA1 (matching the forward strand of the gene): GAACGGCCATTGTCGGTGGCAGG and gRNA2 (matching the forward strand of the gene): CTTCTATGCTAATAACGGGTGGG were selected. These two targeting vectors were used for embryo injections, together with Cas9 mRNA. gRNA generated by *in vitro* transcription was then co-injected into fertilized eggs for the generation of KO mice.

**Figure 1.**
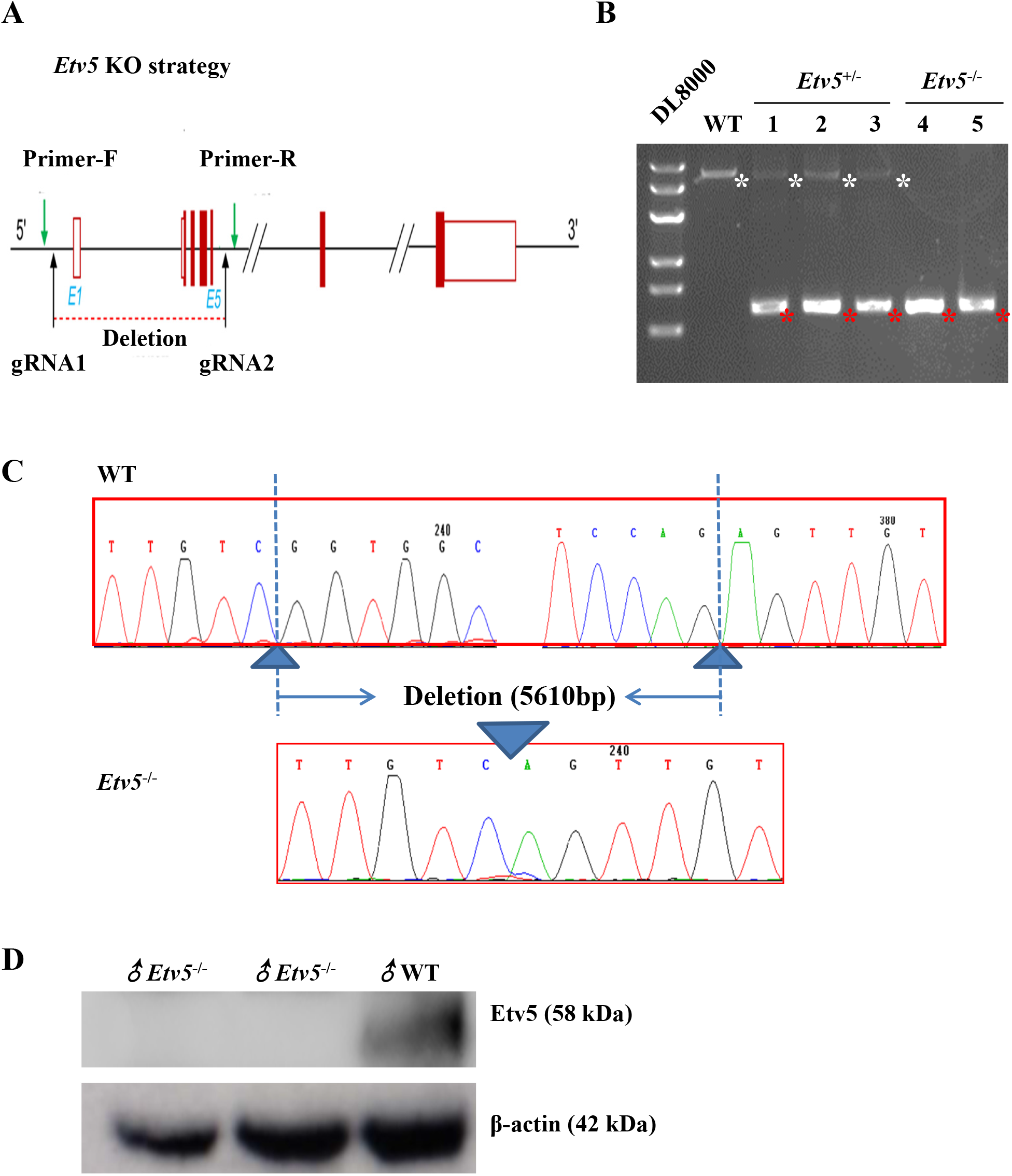
Generation of *Etv5*^*-/-*^ mice by CRISPR/Cas9. (A) Schematic depicting the strategy used for the generation of *Etv5*^-/-^ mice, which included gRNA1 and gRNA2 targeting sites. Primer-F and primer-R were used to genotype the mice. (B) Genotype determined by PCR for newborn male mice littermates. WT, *Etv5*^+/-^, and *Etv5*^-/-^ represent the wild type, heterozygote, and homozygote, respectively. White asterisks indicate WT *Etv5* genomic sequence (6326 bp), and red asterisks indicate the truncated *Etv5* sequence after CRISPR editing (710 bp). (C) Sequencing of the PCR products of *Etv5*^-/-^ mice and WT controls shown in B. (D) Etv5 protein expression in WT and *Etv5*^-/-^ mice measured by Western blot.

### *Etv5* genotyping and expression

For CRISPR-induced mutation assays, female and male heterozygous edited mice were bred to generate homozygous edited offspring. DNA was extracted from the toes of pups from the same litter at day 2 post-birth for PCR analysis and sequencing. The following two primers were used for genotyping by PCR: Mouse *Etv5* Primer-Forward: 5’-CAACTGGTGCCCTTCCCAGTCT-3’, Mouse *Etv5* Primer-Reverse: 5’-GCCGCTCTTAAACCTGTTCATTCG-3’. WT mice were designated as the control group, and *Etv5*^*-*/-^ mice comprised the experimental group. Polyclonal anti-ERM/Etv5 antibody (1:100 dilution; ab102010, Abcam, Cambridge, MA) recognizing amino acids 244 of the human ERM/Etv5 was used to measure Etv5 protein expression in *Etv5*^-/-^ male and WT mice by Western blot. The anti-β-actin monoclonal antibody was used as the loading control.

### Hematoxylin-eosin staining and immunohistochemistry

Whole testes were fixed in 4% paraformaldehyde and then paraffin sectioned for hematoxylin-eosin (H.E.) staining and immunohistochemistry. For H.E. staining, paraffin sections were counterstained with hematoxylin and eosin and then observed under an electron microscope to compare spermatogonium between *Etv5*^-/-^ mice and WT littermates. Immunohistochemistry was performed after antigen retrieval with EDTA (pH 9.0), 3% hydrogen peroxide was used for blocking endogenous peroxidase, and 3% BSA was used to block non-specific binding. Sections were incubated with specific antibodies. Anti-promyelocytic leukemia zinc-finger (PLZF) mouse antibody (sc-28319, Santa Cruz, Dallas, TX), a marker of undifferentiated spermatogonia, was used at 1:100 dilutions. HRP-labeled goat anti-mouse IgG diluted at 1:200 was used as the secondary antibody. The sections were then washed and developed using DAB color rendering and nuclear staining. Hematoxylin-stained nucleus that was blue and brownish yellow color indicated positive expression of DAB.

### Real time PCR

Total RNA samples were extracted using Eastep Super Total RNA Extraction Kit (LS1040, Promega, Madison, WI) according to the manufacturer’s instructions. Total RNA was converted to cDNA using PrimeScript RT reagent Kit with gDNA Eraser (RR047A, Takara, Dalian, China). The mRNA expression levels of the *Ccl9, Prm2*, and *Cyp17a1* genes were then measured by quantitative PCR using PowerUp™ SYBR™ Green Master Mix (A25742, Thermo Fisher Scientific, Austin, TX). β*-Actin* served as an internal control. Each gene from control and experimental samples was tested in triplicate. Relative gene expression was calculated using the 2^-ΔΔCT^method.

### Measurement of testosterone concentrations

Blood was collected from mice via the orbital vein and then centrifuged for serum collection. Testosterone concentrations in *Etv5*^-/-^ male and WT control littermates at 12 weeks of age were determined using the mouse testosterone ELISA kit (E05101m, CUSABIO, Wuhan, China). In brief, 50 μL of standards or samples were added to each well, with blank wells left empty. About 50 μL of HRP conjugate was added to each well except for the blank well. Subsequently, 50 μL of antibody was added to each well. The wells were mixed and then incubated for 1 h at 37 °C. Contents from each well were aspirated and washed three times. The assay plate was blotted dry, and 50 μL of substrate A and 50 μL of substrate B were added to each well and incubated for 15 min at 37 °C in the dark. After incubation, 50 μL of stop solution was added to each well, and the optical density (OD) of each well was recorded within 10 min using a microplate reader set to 450 nm.

### Semen collection and analysis

Mice were euthanized by cervical dislocation. The unilateral epididymis was surgically removed, cut into pieces, and immersed in 1 mL of SpermRinse™ (Vitrolife, Göteborg, Sweden). After incubation at room temperature, 10 μL of sperm solution was transferred to a hemocytometer to determine the number and vitality of sperm. The average of three data records from different mice in each group was then determined. The sperm smear test was used to compare the sperm concentrations between *Etv5*^-/-^ male and WT mice. In brief, approximately 1 mL of sperm solution was centrifuged at 2,000 rpm for 10 min. The resulting precipitate was re-suspended with 500 μL of SpermRinse™, followed by the addition of trypan blue (1:2 in volume). Slides were sealed using glycerol gelatin after 10 μL of the sample was placed onto glass slides.

### SSC preparation

Testes were harvested from 6 to 8 days postpartum C57BL/6 male pups and then digested using a two-step enzymatic digestion protocol. In brief, testes were washed in DPBS with 2% penicillin–streptomycin, and the tunica albuginea and convoluted epididymis were removed. The seminiferous tubules were digested in 5 mL of DPBS solution I consisting of 1 mg/mL collagenase type IV (17104019, Gibco, Grand Island, NY) and 20 U/mL DNase I (2212, Takara, Dalian, China) at 37 °C for 10 min with intermittent agitation every 5 min. Samples were then centrifuged at 300 g for 5 min, and the supernatants were discarded. The precipitates were washed in DPBS and incubated in 5 mL of solution II consisting of 20 U/mL DNase I and 0.25% trypsin/EDTA at 37 °C for 5 min until cells were completely dispersed. Digestion was terminated using 10% fetal bovine serum in DMEM/F12 (11320082, Gibco, Grand Island, NY). The samples containing single cells and clumps were filtered through a nylon cell strainer with 40 μm pore size. Single-cell suspensions were collected in the filtrate and then centrifuged. Supernatants were discarded, and the cell pellet was washed three times with DPBS and then resuspended in complete medium. The number and viability of the resulting dissociated single cells had a density of 1.06×10^7^ cells/mL with viability greater than 99%.

### Lentivirus infection

SSCs were infected with LPP-EGFP-Lv156-400 lentivirus expressing EGFP prior to transplantation to generate *EGFP*-transgenic SSCs. In brief, 30 μL of lentivirus (1×10^8^ TU/mL) was added into 1 mL of cell suspension and then seeded into 12-well plates. The cells were incubated at 37 °C and 5% CO_2_ atmosphere for 12 h to allow lentivirus infection and *EGFP* transgene incorporation in SSC genome. Subsequently, cells were washed once in DPBS and re-suspended gently in DPBS using a Pasteur pipette. Cell suspensions were then transferred into a new 15 mL centrifuge tube and centrifuged. Supernatants were discarded and cell pellet were re-suspended in 300 μL of Hanks’ Balanced Salt Solution (HBSS, 14170112, Gibco, Grand Island, NY). The cell suspension was transferred to a designated transplantation suite for transplantation.

### SSC transplantation

Mice with homozygous mutations of the *Etv5* gene were used as recipients for transplantation. Every recipient was transplanted into only one side of the testis, with the other side of the testis used as the non-transplanted control. Mice were anesthetized by an intraperitoneal injection of anesthetic. An appropriate amount of 1.25% 2,2,2-tribromoethanol (M2910, Easycheck, Nanjing, China, 0.2 mL/10 g body weight) was absorbed in a 1 mL sterile syringe. The needle tip of the syringe was faced upward and then pierced the abdominal cavity at a 45° angle. The drug was slowly injected when the tip part could be moved handily. The mouse’s toe or paw was stimulated, and waiting for it to faint but could keep breathing steadily. At this moment, the mouse was placed dorsally under a stereomicroscope for the transplantation procedure.

The cuticular layer at the midline of the abdomen, not far from the genitals, was lifted using small forceps. A transverse 0.3 cm incision was made, followed by an incision to the peritoneum and abdominal muscle layer until the peritoneal cavity was visible. The lateral fat pad attached to the testis was gently pulled until the testis was exteriorized. The efferent duct that connected the testis to the epididymis was identified, and fat tissues around the duct were gently removed. The SSC suspension was then carefully transferred to a 100 µL volume microinjection syringe. The syringe was connected into a capillary glass tube with an inner diameter of 40–50 µm at the tip. The cell suspension was gently forced into the seminiferous tubules of the testis via the efferent duct by applying pressure to the syringe. The injection pipette was held parallel to the ordinate axis of the efferent duct. The injection rate and cell suspension flow rate were controlled manually by monitoring the movement of the cell suspension in the tubules. Approximately 5–10 μL of the donor cell suspension was injected into each recipient testis. The other testis was not surgically manipulated and served as a control. After transplantation, the testis was placed back into the peritoneal cavity. The incision was closed with one stitch of a 7-0 absorbable suture from the inside out, with the abdominal muscle layer first, followed by the peritoneum layer, and finally the cuticular layer. Mice were returned to their cages, monitored for distress, and assessed regularly until testis samples were collected.

### Identification of allogeneic SSCs

Testes of recipient mice were harvested 2 months after transplantation to determine the functional recovery of spermatogenesis. Recipient testes were fixed in 4% paraformaldehyde and embedded in paraffin wax for histomorphology and immunohistochemistry analyses. Heterologous spermatogenesis was observed under a fluorescence stereomicroscope by using EGFP-positive sperm obtained from the epididymis of recipient mice. Presence of exogenous genes including *Etv5* and *EGFP* in the testes and spermatozoa were measured by PCR.

### Statistical analysis

PASW Statistics 21 (IBM SPSS, Chicago, IL) was used to determine statistical significance and standard deviation. Body weight, testis weight, gene expression level, testosterone concentrations, and cell counts between WT and KO groups were analyzed using two-tailed unpaired *t*-test. Differences were considered significant at *P*<0.05 (*) and *P*<0.01(**).

## Results

### Generation of *Etv5*^*-/-*^ mice by using CRISPR/Cas9

We designed two gRNAs targeting the introns on both sides of exon 1 (gRNA1) to exon 5 (gRNA2) to delete a 5610 bp fragment of the *Etv5* gene to generate *Etv5*^*-/-*^ mice by embryo injection of Cas9 mRNA and gRNAs (**Fig. 1A**). The efficiency of CRISPR-induced founder mice is shown in **Table 1**. The genotypes of the founder and their offspring were identified using primers F/R (**Fig. 1A**). DNA gel electrophoresis results demonstrated that homozygous KO mice generated a 710 bp band, heterozygotes had two bands (710 and 6326 bp), and wild-type mice had a single band of 6326 bp (**Fig. 1B**). PCR and DNA sequencing demonstrated that homozygous KO mice had a 5610 bp deletion (**Fig. 1C**). In addition, Western blot analysis confirmed a lack of Etv5 expression in homozygous *Etv5*^*-/-*^ mice (**Fig. 1D**). Interestingly, *Etv5*^-/-^ male mice at 3 weeks of age were obviously smaller than WT male mice, and this difference was more pronounced at 12 weeks of age (**Supplementary Fig. 1A**). The body weights of *Etv5*^-/-^ male mice were markedly lower at each point from 14 postnatal days to 84 days (*P*<0.05) compared with WT controls, with the magnitude of body weight decline between 34.3% and 45.7% (**Supplementary Fig. 1B**).

**Table 1.** Generation of CRISPR-induced founder mice with *Etv5* deletion.

| **Breed** | **Experimental dates** | **Injected embryos** | **Surrogates** | **Pregnants** | **Founders** | **Positive founders<sup>1</sup>** |
| --- | --- | --- | --- | --- | --- | --- |
| B6N*B6N | June. 2. 2017 | 128 | 4 | 3 | 12 | 1 |
| B6N*B6N | June. 2. 2017 | 141 | 5 | 2 | 7 | 0 |
| B6N*B6N | July. 24. 2017 | 179 | 7 | 4 | 37 | 1 |

<sup>1</sup> Identified by PCR showing a truncated *Etv5* gene

### Lack of SSCs in the seminiferous tubules of *Etv5*^*-/-*^ mice

We measured the weight of testes in *Etv5*^-/-^ mice from day 4 to 12 weeks of age and compared them with that in WT mice of the same age. The weights were almost similar in mice aged before 1 week. However, the testis weights of *Etv5*^-/-^ mice were markedly lower from 3 to 12 weeks of age (*P*<0.01) compared with those of WT controls, with the magnitude of testis weight declining between 43.1% and 67.4% (**Fig. 2A**). The testis-to-body-weight ratios were also significantly lower in *Etv5*^-/-^ mice than in WT mice after 3 weeks of age (**Fig. 2B**). The difference was also reflected in the sizes of testes; the testes of *Etv5*^-/-^ mice were much smaller than those of WT mice at 3 weeks of age, and the difference became more evident at 12 weeks of age (**Fig. 2C**). H.E. staining demonstrated a remarkable decrease in the number and type of testicular cells in the seminiferous tubules of *Etv5*^-/-^ mice at 3 weeks of age. Testicular cells were absent in the tubules of *Etv5*^-/-^ mice, and Sertoli cell-only tubules were observed at 12 weeks of age (**Fig. 2D**). Additionally, we measured the protein expression levels of PLZF, which is a marker for SSCs. We observed that PLZF was still expressed in *Etv5*^*-/-*^ mice in some seminiferous tubules at 3 weeks of age. However, by 12 weeks of age, SSCs in *Etv5*^-/-^ had completely disappeared but were present in WT mice (**Fig. 2E**). In addition, the numbers of premeiotic germ cells, meiotic germ cells, and round spermatids significantly declined in *Etv5*^*-/-*^ mice and were undetectable by 12 weeks of age (**Fig. 2F**). Additionally, the percentage of vacuolation observed in seminiferous tubules in both 3- and 12-week-old *Etv5*^-/-^ mice were higher than that in WT control mice, reaching 100% at 12 weeks of age (**Fig. 2G**). The diameter of seminiferous tubules of *Etv5*^-/-^ mice was significantly smaller than that of WT controls at both 3 and 12 weeks of age (*P*<0.01; **Fig. 2H**).

**Figure 2.**
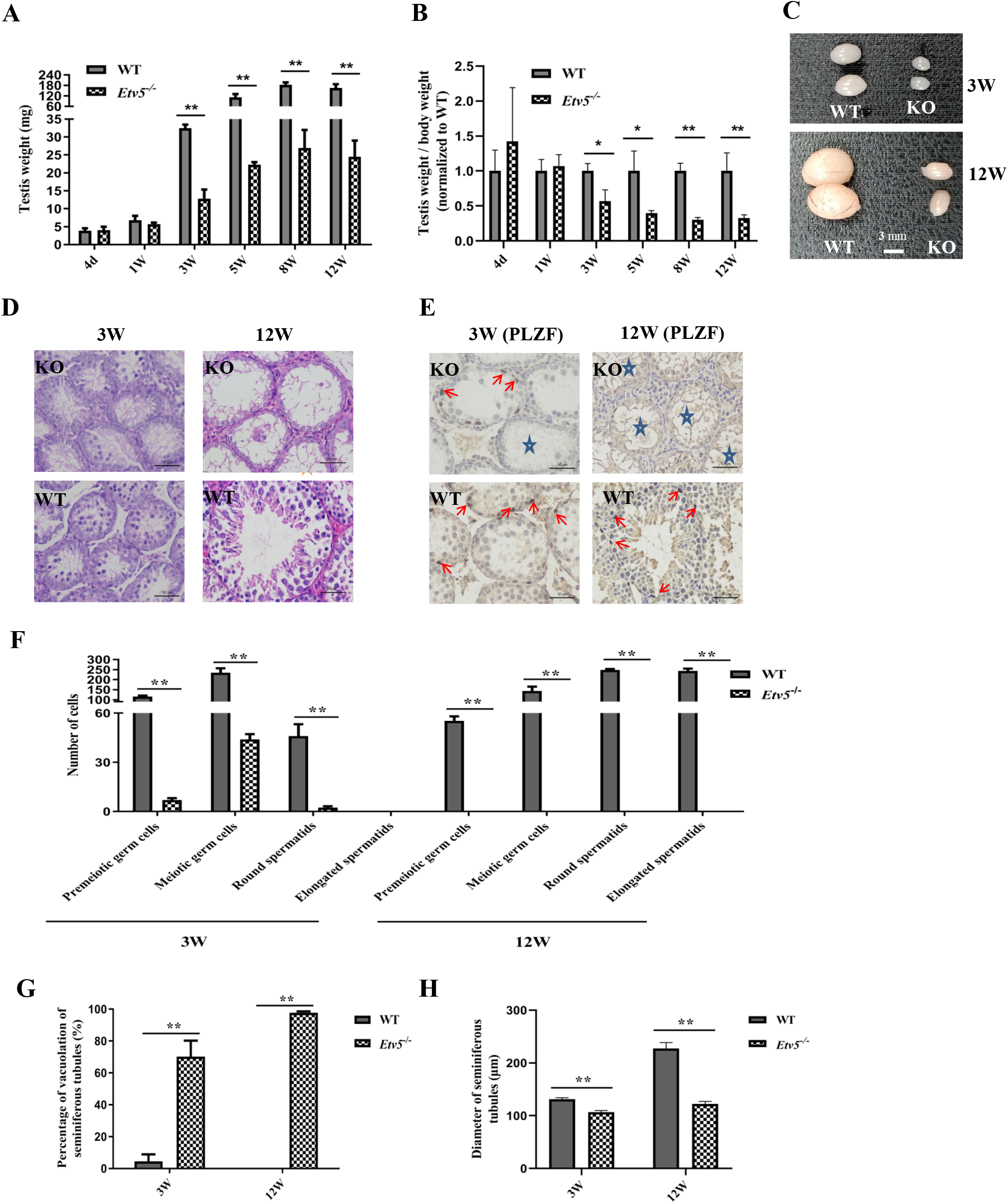
Absence of SSCs in the seminiferous tubules of *Etv5*^*-/-*^ mice. (A) Weight of testes in *Etv5*^-/-^ mice compared with WT littermates at periodic points during development. For all time points, n = 4 per group for both WT and *Etv5*^-/-^ mice. (B) Testis-to-body-weight ratio of *Etv5*^-/-^ mice and WT littermates (n = 4). (C) Comparison of size of testes of WT and *Etv5*^*-*/-^ mice at 3 and 12 weeks of age. Scale bars = 3 mm (D and E) Representative images for H.E. staining and PLZF immunohistochemistry of testes in WT and *Etv5*^-/-^ mice littermates at 3 and 12 weeks. Red arrows indicate SSCs with brown positive staining by PLZF antibody. Pentacles reveal vacuolation in some seminiferous tubules in *Etv5*^-/-^ mice. Scale bars = 50 μm. (F) The numbers of premeiotic germ cells, meiotic germ cells, round spermatids, and elongated spermatids in seminiferous tubules in WT and *Etv5*^-/-^ mice littermates at 3 and 12 weeks. The cell numbers were counted from the cross sections of seminiferous tubules in a 200× visual field. Samples from 3 mice were prepared, and data were collected from 3 slices per mice. (G) The percentage of vacuolation of seminiferous tubules. (H) Diameter of seminiferous tubules from WT and *Etv5*^-/-^ mice at 3 and 12 weeks of age. n = 3 per group shown in F, G, and H for both WT and *Etv5*^-/-^ mice. Nine slices (3 for each mouse) per group were prepared, and data from each slice were collected from at least three different regions. Data are means ± standard deviation (S.D). * represents *P*<0.05, ** represents *P*<0.01, determined by unpaired *t*-test.

We next examined sperm production in KO mice at different ages. The results were consistent with the findings of H.E. staining and immunohistochemistry. At the time of sexual maturity (6 weeks of age), there were numerous motile sperm in WT mice compared with age-matched *Etv5*^-/-^ mice, whose sperm numbers were extremely low (**Table 2** and **Supplementary Fig. 2**). At 12 weeks of age, numerous motile sperm were observed in the epididymis WT mice, but there was a lack of spermatozoa in age-matched *Etv5*^-/-^ mice (**Table 2** and **Supplementary Fig. 2**). In addition, we found that the expression levels of the *Etv5* target gene *Ccl9*, spermatid-specific gene *Prm2*, and interstitial gland-specific gene *Cyp17a1* were significantly decreased (*P*<0.01, *P*<0.01, and *P*<0.05, respectively) in *Etv5*^-/-^ mice (**Supplementary Figs. 3A, 3B, and 3C**). We also tested testosterone concentrations in WT and *Etv5*^-/-^ littermates at 6, 8, and 12 weeks. A significant reduction in testosterone concentrations was observed in *Etv5*^-/-^ mice compared with WT mice, and a maximum difference was reached at 12 weeks of age (*P<*0.01; **Supplementary Fig. 3D**).

**Table 2.**
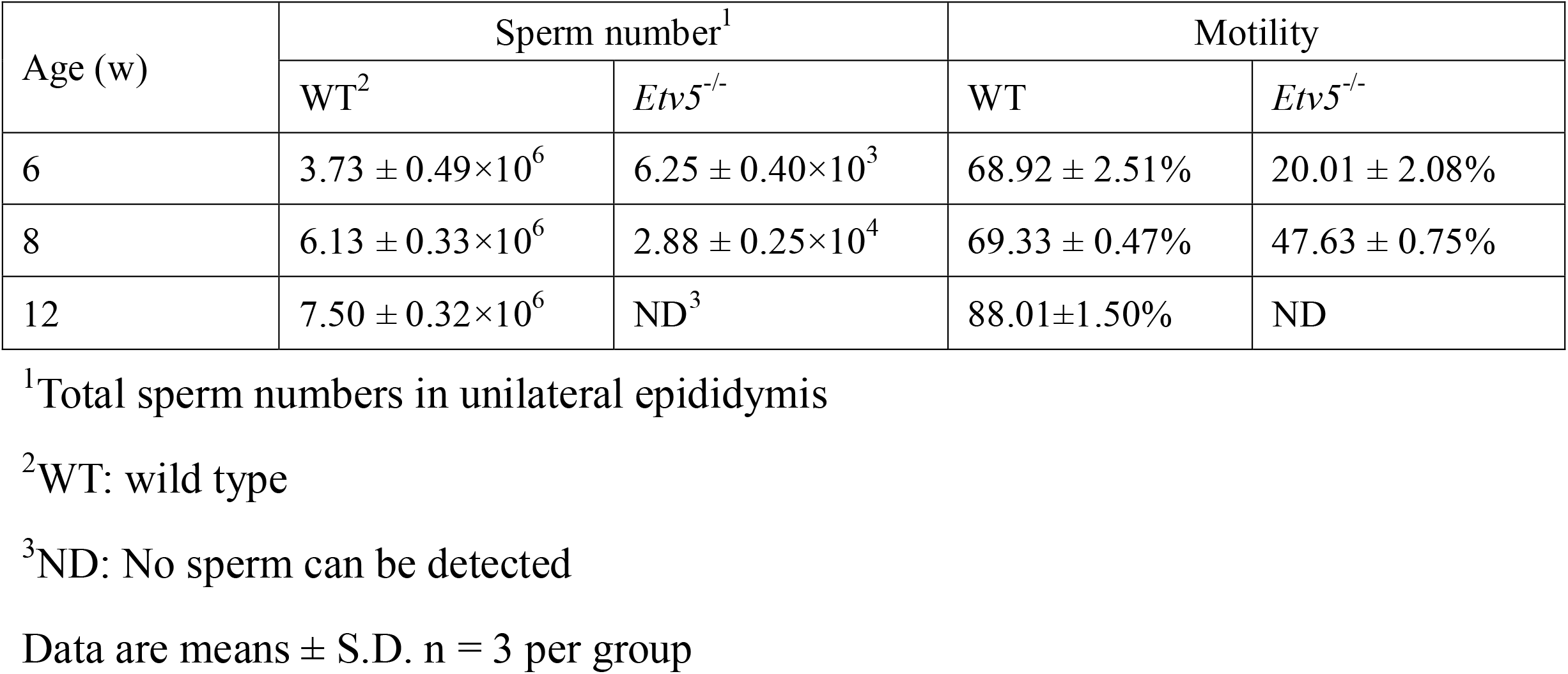
Number and motility of sperm in one side epididymis between WT and *Etv5*^-/-^ mice.

**Figure 3.**
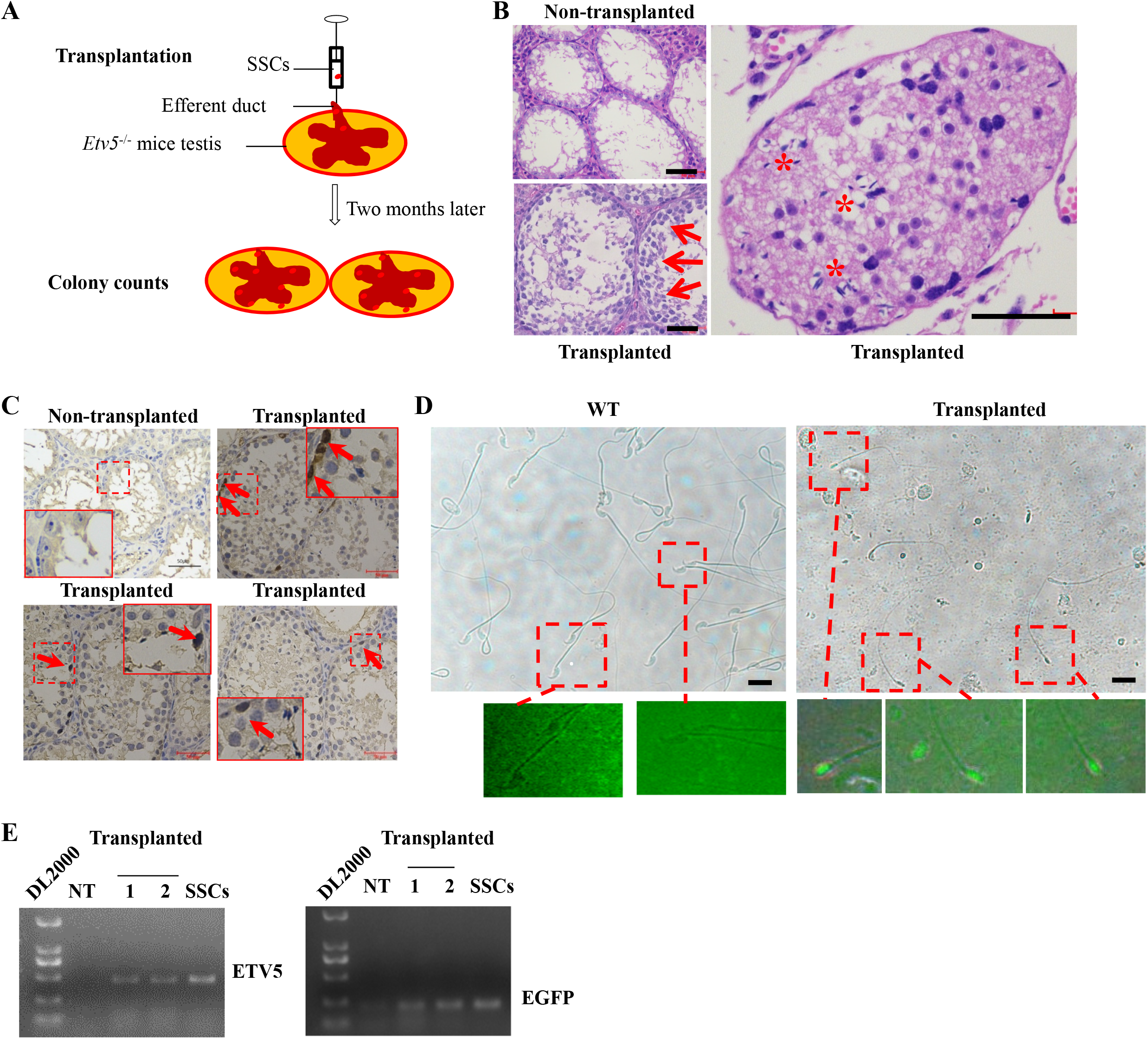
Transplantation of allogeneic SSCs through the efferent duct. (A) Transplantation process of allogeneic SSCs through the efferent duct. (B) H.E. staining of non-transplanted and transplanted testes. Red arrows indicate restored germ cell layers in the seminiferous tubules, and red stars indicate regenerated immature sperm. Scale bars = 50 μm. (C) PLZF immunohistochemistry of testes showing restored spermatogonia with brown positive staining (red arrows) in the seminiferous tubules. The insets represent high magnification of pointed cells in break line boxed areas. Scale bars = 50 μm. (D) Sperm collected from epididymis 2 months post-transplantation and age and breed-matched WT mice. Red boxes point to regenerated sperm with EGFP expression in *Etv5*^-/-^ mice and WT sperm which are EGFP negative in the same conditions. Scale bars = 20 μm. (E) heterologous *Etv5* and *EGFP* DNA were measured to determine the origin of epididymal sperm. NT, non-transplanted testes; 1, transplanted testes; 2, sperm collected from epididymis of SSC-transplanted testes; SSCs, donor cells used for transplantation.

### Transplantation of allogeneic SSCs through the efferent duct

To determine whether the *Etv5*^-/-^ mouse can serve as a recipient model for SSC transplantation, we transplanted allogeneic SSCs (expressing EGFP by lentiviral transduction) through the efferent duct into immature (3 weeks of age) and mature (7 weeks of age) mice (**Fig. 3A** and **Table 3**). Our preliminary experiments showed that almost all lentiviral transduced SSCs were positive for EGFP expression at a multiplicity of infection of 5:1 (viruses to cells). In nine immature recipients, seven harbored spermatozoa in epididymis at 2 months post-transplantation (**Table 3** and **Supplementary Fig 4**). The average sperm number of the seven restored mice was 9.58 ± 5.05×10^4^ (**Table 3**). Morphological analysis of the transplanted testes found that the proportion of tubules with restored spermatogonia was 29.51 ± 0.08% in the seven restored testes (**Table 3**). Reconstruction of germ cell layers (**Fig. 3B**, left lower) and regeneration of elongated spermatozoa (**Fig. 3B**, right) could be observed in these tubules, and spermatogonia were positive for PLZF protein expression (**Fig. 3C**, transplanted groups) in *Etv5*^-/-^ mice after SSC transplantation. No spermatozoa and germ cells were detected in non-transplanted immature testes (**Figs. 3B** and **3C**). To determine the origin of the sperm from post-transplanted immature testes, we examined them under a fluorescence microscope. The regenerated sperm emitted green light, indicating they were of donor SSC origin (**Figs. 3D**). Furthermore, the presence of heterologous genes *Etv5* and *EGFP* was measured in recipient epididymal sperm by PCR. *Etv5* and *EGFP* were not detected in non-transplanted testes; however, they were present in spermatozoa of transplanted *Etv5*^-/-^ mice (**Fig. 3E**). This finding further indicated that the sperm were derived from donor cells. In addition, no spermatozoa were detected in all nine mature recipients with SSC transplantation (**Table 3**).

**Table 3.** Development of allogeneic spermatogenesis in *Etv5*^-/-^ recipient mice after 2 months.

| Recipient mice<sup>1</sup> |  | Non-transplanted testes |  |  | Transplanted testes |  |  |  |  |
| --- | --- | --- | --- | --- | --- | --- | --- | --- | --- |
| Age (w) | number | Testes | Tubules | Epididymides | Testes<sup>3</sup> | Tubules<sup>4</sup> | Epididymides<sup>5</sup> | Sperm<sup>6</sup> | Motility |
| 3 | 9 | ND<sup>2</sup> | ND | ND | 9/9 (100%) | 29.51 ± 0.08 | 7/9 (77.78%) | 9.58 ± 5.05×10<sup>4</sup> | 0 |
| 7 | 9 | ND | ND | ND | ND | ND | ND | ND | ND |

<sup>1</sup>Every recipient with one side testis transplanted and the other side testis untransplanted
<sup>2</sup>ND: No sperm can be detected
<sup>2</sup>Testes with sperms
<sup>4</sup>Percentage of seminiferous tubules with repopulated germ cells
<sup>5</sup>Epididymides with sperms
<sup>6</sup>Average sperm numbers in epididymides of 7 mice with restored *spermatogenesis*
Data are means ± S.D.

## Discussion

Etv5 is essential for SSC self-renewal and KO of Etv5 severely impairs SSC development and results in male infertility (Chen., 2005; Hofmann., 2008; and Ishii., 2012). Chen et al. demonstrated that *ERM*^*-/-*^ (*Etv5*^*-/-*^) mice undergo a progressive germ cell depletion with a gradual loss of spermatogonia in the seminiferous tubules staring from 4 weeks of age. *ERM*^*-/-*^ tubules lost most germ cells, but containing morphologically normal Sertoli cells at the basement membrane by 10 weeks of age (Chen., 2005). In our study, we successfully generated *Etv5*^*-/-*^ mice by using CRISPR/Cas9 system for embryo injection. The generated KO mice gradually lost SSCs but preserved an intact structure of seminiferous tubules with a similar time frame as previous report, indicating that they could be potential germ cell-free models for SSC transplantation.

Our KO manipulation revealed that undifferentiated and differentiated spermatogonia were lost in the majority of seminiferous tubules, and only a small region of the tubules retained multilayer germ cells in immature mice. *Etv5*^-/-^ mice at 3 weeks old had partial germ cell loss in the seminiferous tubules and their testicular sizes were slightly smaller than those of age-matched WT controls. By 12 weeks of age, the testicular sizes of *Etv5*^-/-^ mice were much smaller than those of WT controls, with a severe germ cell lost in the tubules. Interestingly, 3-week-old *Etv5*^-/-^ mice were smaller in body size and weight compared with WT mice. At 12 weeks of age, this trend was even more obvious. The reduced body weights in *Etv5*^-/-^ mice indicated that *Etv5* had an influence on overall growth. *Etv5* mRNA has been detected in a variety of tissues, including the heart, lungs, thymus, lymphocytes, kidneys, and skeletal muscles (Liu., 2003; T’Sas., 2005). The widespread expression of *Etv5* during development may be crucial for growth. In a previous study that investigated the viability of *Etv5*^*-/-*^ mice, *Etv5* mRNA was found to be abundantly expressed in the brain, lungs, and colon, but it was most abundantly expressed in the testes (Schlesser., 2008).

We next investigated whether our *Etv5*^*-/-*^ mouse model would support donor-derived spermatogenesis and sperm generation after allogeneic SSC transplantation. Previous report demonstrated that *Etv5*^*-/-*^mice showed serious defects in Sertoli cells, which could not form an optimal testicular environment to support spermatogenesis (Chen., 2005). However, the overall structure of seminiferous tubule remains intact in spite of severe loss of germ cells and other somatic cells. Also, testicular cell preparation and transplantation process is usually accompanied by existence of lots of testicular somatic cells, which could supplement the defects of Sertoli cells in *Etv5*^*-/-*^ mice. In the present study, we definitely obtained donor-derived spermatozoa in transplanted *Etv5*^*-/-*^ mice. Although spermatogenesis was visible in transplanted immature *Etv5*^*-/-*^ mouse testes, the number and motility of regenerated sperm in the epididymis were not ideal. We guess multiple reasons could contribute to the inefficiency in SSCT transplantion in *Etv5*^*-/-*^ models, including immunological rejection between donor and recipient, impaired Sertoli cells constitution or stem cell microenvironment by absence of Etv5, or inadequate quality and number of implanted SSCs. Also, recoverable spermatogenesis following SSC transplantation seems to be slower than endogenous spermatogenesis. Complete spermatogenesis takes approximately 34.5 days from type A single (As) spermatogonia to mature spermatozoa in WT mice (Oakberg., 1956). Transplanted SSCs usually need 2 weeks for a complete colonization. Meiosis is usually initiated within the second month, with several spermatids being observed after 2-month transplantation. The degree of germ cell differentiation will continue to increase, with normal spermatogenesis being observed 3 months post-transplantation (Nagano., 1999). Therefore, the time point of 2 months in our experiment could be not enough for detection of a full round of spermatogenesis following SSC transplantation.

Our results showed SSC transplantation effect between immature and mature *Etv5*^*-/-*^ testis is significantly different. SSC transplantation in immature *Etv5*^*-/-*^ testis is more feasible than mature *Etv5*^*-/-*^ testis, which is in line with previous studies which showed that SSC transplantation in immature pup testes is more efficient than adult testis (Shinohara.,2001; Ishii.,2013). As a lack a fully formed blood-testis barrier, transplantation of testis cells into pup testis has a 5-to 10-fold increased colonization efficiency compared with mature testis. The area for colonization per donor stem cell is also 4 times larger in recipient pups than adults (Shinohara., 2001). These factors facilitate a more efficient restoration of fertility by SSC transplantation in infertile immature recipients. Furthermore, immature *Etv5*^*-/-*^ testis at the age of 3 weeks harbors residual endogenous spermatogonia which could facilitate maintenance of testicular function and help restoration of fertility after transplantation (Kanatsu-Shinohara., 2016).

In summary, we generated *Etv5*^-/-^ mice with a genetic ablation of germline. SSC donors transplanted into recipient testes could partially restore heterologous spermatogenesis and produce donor-derived sperm, even though their quantity and vitality were not optimal. SSC transplantation in this genetically modified mouse models is possible but remains in a low efficiency in the present report. Further investigations are required to optimize the SSC transplantation process in the modified models to obtain ideal transplantation outcomes.

## Acknowledgment

This work was supported by the National Natural Science Foundation of China (31772555).

## Author Contributions

Conceptualization: Huaqiang Yang, Zhenfang Wu, and Bin Chen; Investigation: Xianyu Zhang, Mao Zhang, Xin Zhao, and Pingping Xing; Formal analysis: Huaqiang Yang, Xianyu Zhang, Guoling Li, and Zicong Li; Funding acquisition: Huaqiang Yang and Zhenfang Wu; Drafting and revision of manuscript: Xianyu Zhang and Huaqiang Yang. All authors contributed to the manuscript. The authors declare no competing interests.

## Figure legends

**Supplementary Fig. 1. Body weight of KO mice**. (A) The phenotype of WT and *Etv5*^-/-^ mice at 3 and 12 weeks of age. Arrows indicate *Etv5*^-/-^ and WT mice. (B) Body weight of WT and *Etv5*^-/-^ mice at periodic time points during development from 4 postnatal days to adulthood (12 weeks). For all time points, n = 5 for both WT and *Etv5*^-/-^ mice. Data presented as mean ± S.D. \**P*<0.05, \*\**P*<0.01, assessed by unpaired *t*-test.

**Supplementary Fig. 2. Sperm density in the epididymis in 6-, 8-, and 12-week WT and *Etv5***^**-/-**^ **mice**. Arrows indicate sperm and the insert represents the higher magnification of sperm. Scale bars = 50 μm.

**Supplementary Fig. 3. Changes in gene expression related to spermatogenesis and testosterone concentrations**. mRNA expression levels of *Etv5* target gene *Ccl9* (A), spermatids-specific gene *Prm2* (B) and interstitial gland-specific gene *Cyp17a1* (C) in the testes of WT and *Etv5*^-/-^ mice littermates at 12 weeks. WT values were set as 100%. (D) Testosterone concentrations of WT and *Etv5*^-/-^ littermates at 6, 8 and 12 weeks. n = 3 per group. Data presented as mean ± S.D. \**P*<0.05, \*\**P*<0.01, assessed by unpaired *t*-test.

**Supplementary fig. 4. Sperm regeneration**. The sperm collected from epididymis in recipient mice 2 months post-transplantation. Mature sperm with intact shape can be observed (Red arrows). Scale bars = 50 μm.

