## Supplementary figures and images for "Establishment of *Etv5* gene knockout mice as a recipient model for spermatogonial stem cell transplantation"

### Supplemental Figures

A

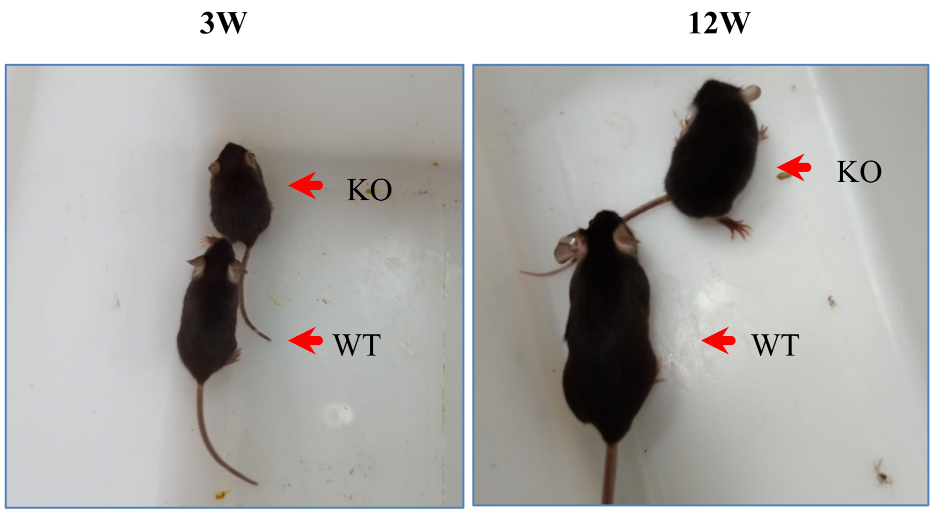


B

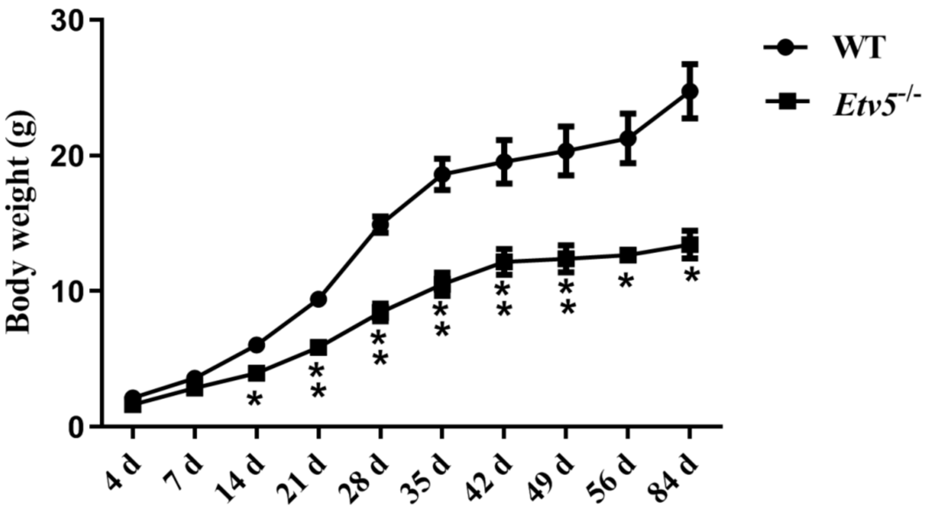


Supplementary Fig. 1

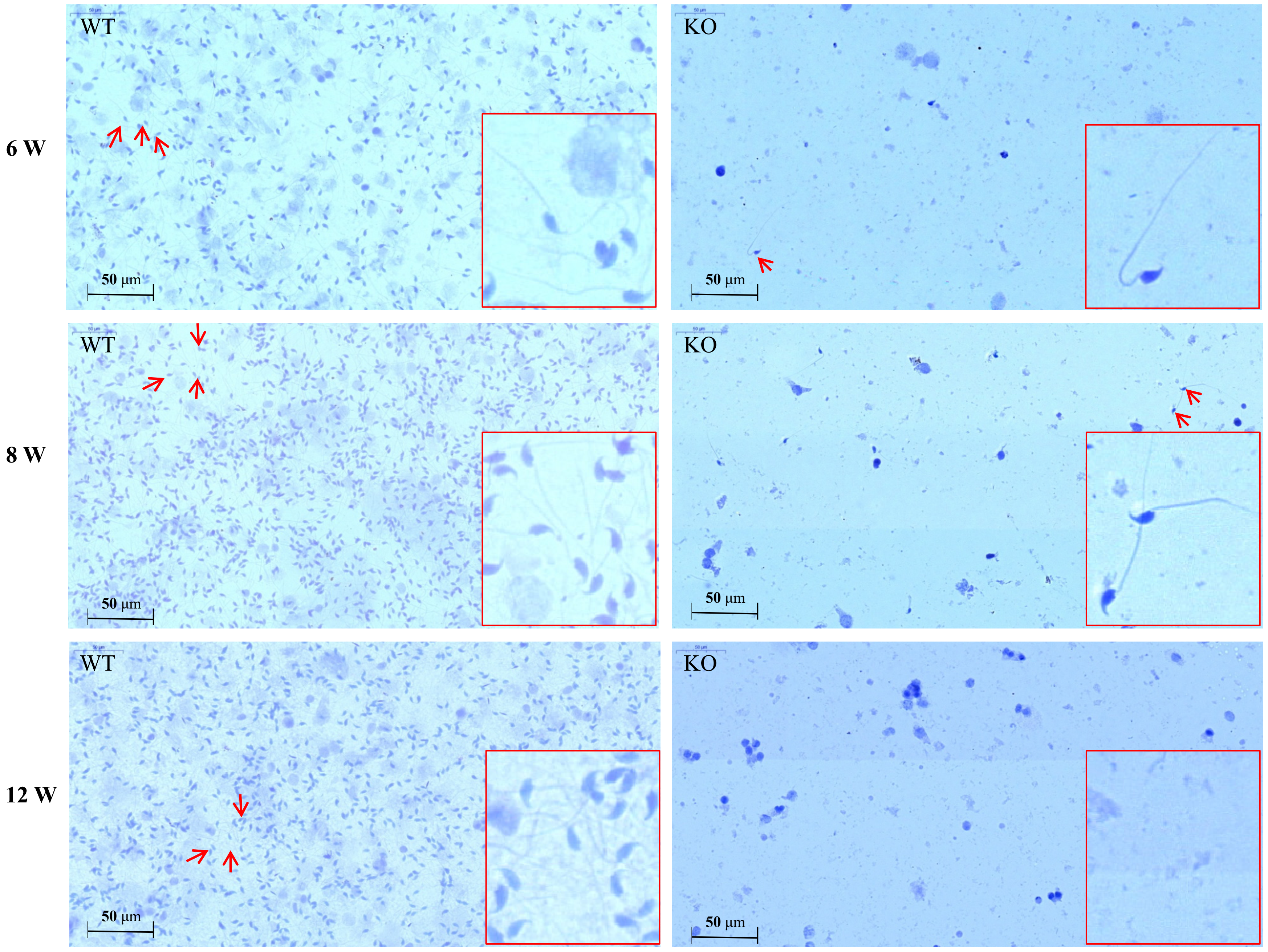


**Supplementary Fig. 2**

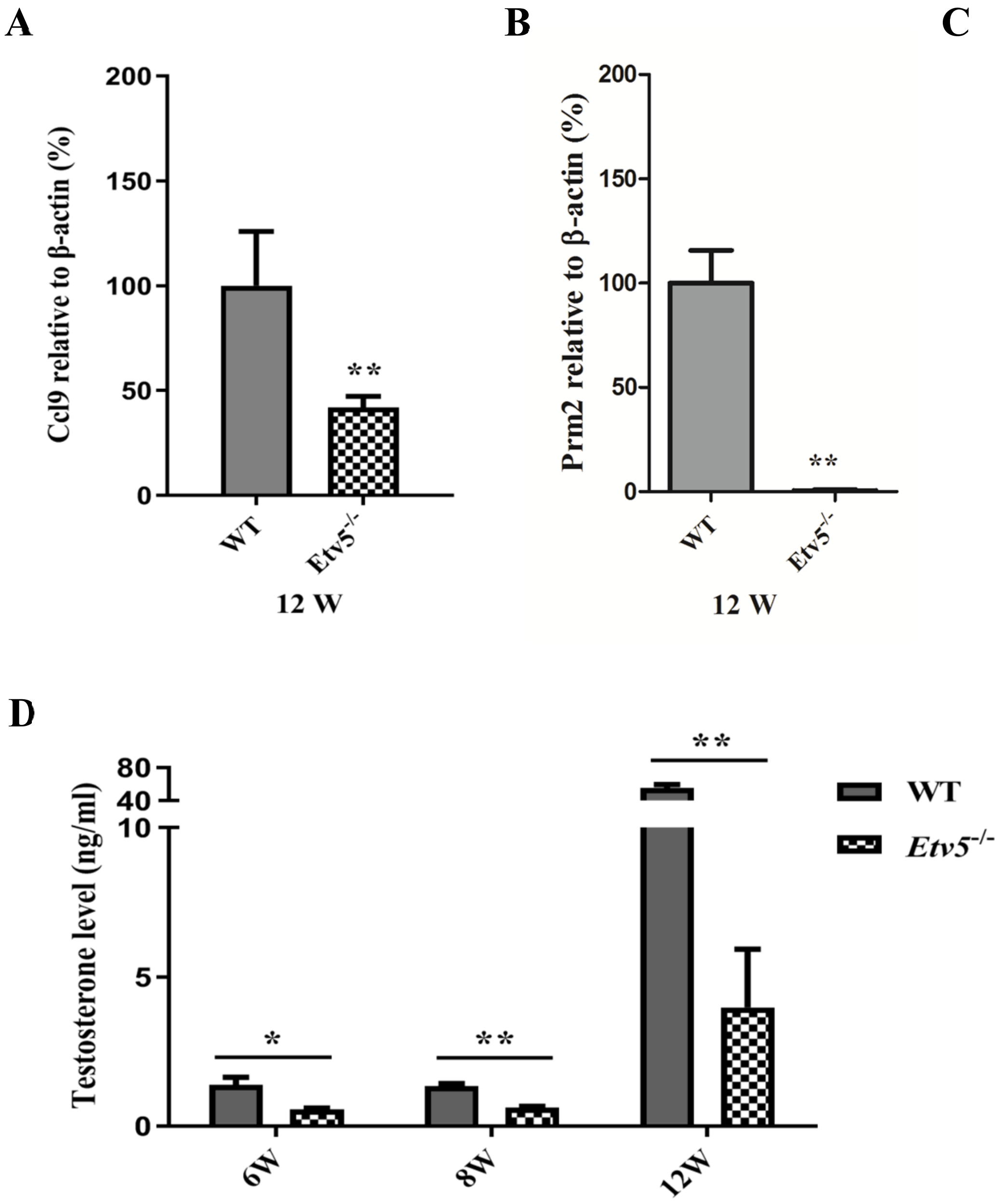


Supplementary Fig. 3

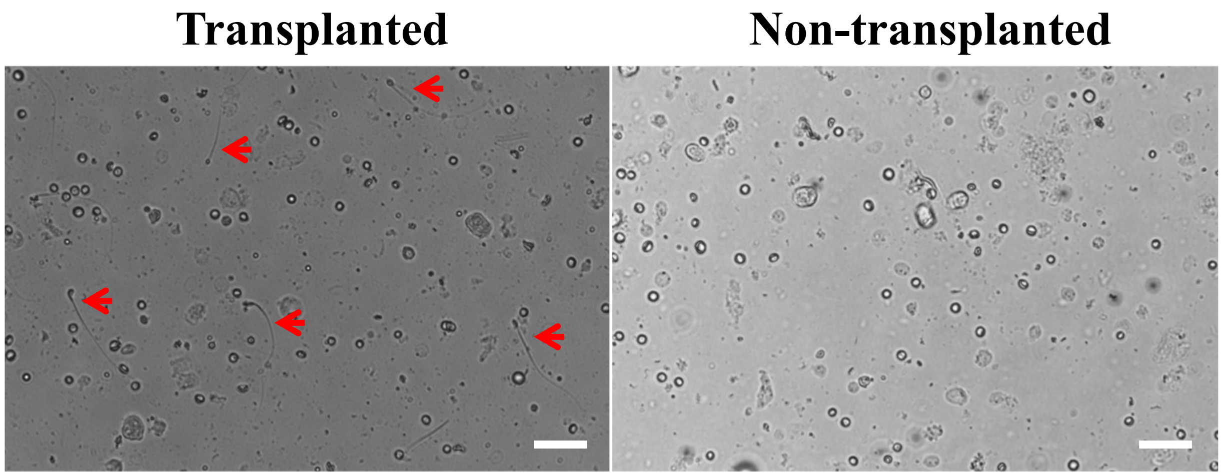


Supplementary Fig. 4
